# Computational analysis of the plasmodium falciparum-induced intraerythrocyte deformations: insights into malaria parasite egress mechanics and eryptosis

**DOI:** 10.1101/2024.05.17.594756

**Authors:** Chimwemwe Msosa, YD Motchon

## Abstract

In malaria infection, early eryptosis eliminates the protective niche provided by the infected erythrocyte, thus potentially interfering with the parasite’s survival rate in the human host, consequently presenting a putative target mechanism for antimalarial therapeutic interventions. The malaria parasite-induced oxidative stress is the primary trigger of eryptosis. Despite being barely investigated, erythrocyte membrane mechanical deformations induced by the malaria parasites during the intraerythrocyte development stage, represent a potential eryptosis trigger.

A finite element model of the plasmodium falciparum-infected erythrocyte was developed and calibrated in Abaqus using pre-determined optical tweezer data of the trophozoite-infected erythrocyte. The developed model computationally predicts mechanistic correlations between erythrocyte membrane areal strain, eryptosis, erythrocyte membrane shear modulus and the volume fraction of malaria parasites in the infected erythrocyte.

The model predicts the erythrocyte membrane areal strain of 3.1 % at the established rupture volume fraction (VF) of 83%, which falls within the pre-determined erythrocyte membrane lysis threshold of 2-4 %. When the erythrocyte membrane in-plane shear modulus is increased from 2.84 µN/m to 131 µN/m, the erythrocyte areal strain increases from 1.55 % to 3.2 % at the same rupture volume fraction (VF) of 83% implying that increasing the erythrocyte membrane stiffness during the malaria intra-erythrocytic development stage can potentially induce early lysis while decreasing the erythrocyte membrane stiffness can potentially induce late lysis.

Understanding the mechanisms governing the exit of malaria parasites from infected erythrocytes during the late schizont stage is crucial for developing effective therapeutic interventions. Existing studies lack a comprehensive exploration of how malaria parasite-induced remodelling affects the areal strain of the erythrocyte membrane. Experimental challenges in studying infected erythrocytes have limited progress, making computational models a valuable tool. This research provides valuable insights into the mechanics of malaria-induced erythrocyte remodelling, offering a computational framework for studying parasite egress to inform potential therapeutic strategies.

## 1 Introduction

Malaria is a devastating infectious disease, causing millions of deaths in sub-Saharan Africa. In 2016, 91 countries reported a total of 216 million cases of malaria, an increase of 5 million cases over the previous year, with an estimated 445,000 deaths, of which the majority occurred in young children of sub-Saharan Africa [1]. Despite the recent global reduction in malaria cases, the lack of efficacious antimalarial drugs due to the spread of persistent antimalarial drug resistance presents a major challenge to the treatment of malaria [2]. As such, novel alternative treatment options are urgently needed [3], and for this reason, host-directed therapies that interfere with host molecules and pathways have emerged in recent years as a potential and novel approach to tackle malaria infections [4]. Among these strategies include approaches that interfere with host cell regulatory factors that govern eryptosis [5, 6] and are of interest for developing long-lasting and effective antimalarial drugs [7].

The exit of plasmodium falciparum parasites from the infected erythrocyte has clinical implications characterized by high fever, nausea, and severe headache, marking the pathogenesis of complicated and uncomplicated malaria [8]. The infected erythrocyte provides a protective niche that supports the malaria parasite’s survival, facilitating schizogony and, subsequently, the egress of merozoites into the human host bloodstream, thus increasing the parasite’s virulence [9]. Soon after invading the erythrocyte, the malaria merozoite undergoes a morphological transformation from an egg-shaped to disc-shaped geometry with a diameter of 2–3 μm and a thickness of ∼0.5 μm [10]. However, the shapes and sizes of the infected erythrocyte in the ring stage when the merozoite is in disk form are similar to those of a healthy erythrocyte, with a small percentage of the host volume occupied by the parasite [11]. During the trophozoite stage, the parasite is at its metabolically most active state. It secretes and exports proteins onto the erythrocyte membrane that modify its structure and permeability. As a result, the membrane’s elastic modulus increases [12, 13] while the cell’s shape, volume, and surface area also change [11, 14]. Schizogony involves the asynchronous intracellular production of multiple nuclei, culminating in a single mass cytokinesis event. The efficiency of parasite replication is essential in controlling the virulence of malaria disease, which tends to depend on parasite loads [15, 16]. The malaria parasite primarily orchestrates its exit from the infected erythrocyte primarily through eryptosis [6]. Therefore, host factors pivotal in regulating eryptosis represent potential targets for host-driven antipathogen therapeutic interventions [17]. However, it should be noted that promoting or preventing cell death can be either beneficial or detrimental for both the malaria parasites and the human host, and this outcome depends on various intrinsic variables related to the infection, including the nature of the pathogen, pathogen load, infection site, and the timing of cell death during the infection [18].

The malaria parasite-induced eryptosis begins with oxidative stress build-up [19] due to the accumulation of byproducts derived from host haemoglobin digestion, leading to the opening of calcium ion channels and the influx of cytoplasmic calcium, consequently causing the externalization of phosphatidylserine (PS) and subsequently inducing eryptosis [20]. In addition to oxidative stress, mechanical deformation of the erythrocyte membrane represents a potential trigger for eryptosis in malaria infection. While the link between eryptosis and erythrocyte membrane deformation in malaria infection remains unknown, previous studies not involving malaria infection have established the link between erythrocyte lysis and the areal strain of 2-4% with an average value of 3% [21].

The link between eryptosis and erythrocyte membrane deformations during the malaria parasite intra-erythrocyte development requires a mechanistic description that accounts for the evolution of erythrocyte membrane mechanical properties, intra-erythrocyte parasite expansion [15, 16] and infected erythrocyte morphological alterations [22, 23]. Furthermore, parasite-induced mechanical deformation can potentially promote the activation of mechanosensitive calcium channels, facilitating the influx of Ca^2+^ into the cell and subsequently inducing eryptosis[24–26].

Our comprehension of eryptosis regulation remains limited by the absence of a clear understanding of how the increase in infected erythrocyte volume caused by plasmodium schizogony, which alters erythrocyte shape [27, 28], and the malaria parasite-induced changes to the erythrocyte membrane [29] impacts eryptosis. As such, the present study aimed to computationally explore the mechanistic link between eryptosis and the mechanical deformations of the erythrocyte. The first objective of the study was thus to develop and validate the malaria parasite-infected erythrocyte (MPIE) finite element model (FEM) for predicting the mechanics of the infected erythrocyte prior to the exit of malaria parasites. The Developed model was initially validated using previously determined diametric data of the trophozoite stage erythrocyte [30]. The malaria parasite volumetric growth and erythrocyte membrane models were developed and calibrated with previously determined data. The second objective was to establish the link between malaria-induced remodelling and eryptosis by conducting in-silico computational studies utilizing the developed FE model.

In the present study, we computationally show that mechanical deformation due to the growth of malaria parasites is linked to malaria-induced eryptosis. Our results demonstrate that increasing the erythrocyte membrane stiffness during the malaria intra-erythrocytic development stage can potentially induce early lysis. Conversely, severely reducing the erythrocyte membrane stiffness can potentially delay lysis. The new insights from this study are thus instrumental in guiding new strategies and developing interventions for treating malaria.

## 2 Methods and materials

### 2.1 Discoid shape model of the infected erythrocyte

A 3D biconcave geometry of the erythrocyte membrane and cytosol was developed using the Eqn(1) [31].

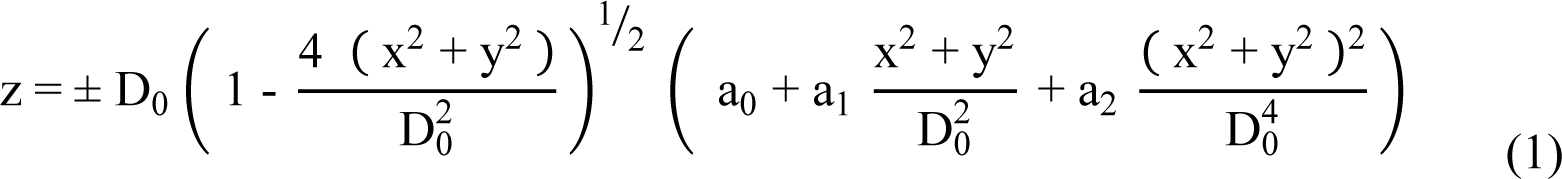

Where x, y, and z are principal coordinate directions, and D_0_ = 7.0 µm is the undeformed diameter of the erythrocyte cell. a_0_, a_1_, and a_2_ are shape parameters with values of 0.132, 2.026, and -4.491, respectively (see Table 1). These shape parameters were manually determined such that the volume and surface area of the trophozoite-infected erythrocyte model were consistent with the literature [31], i.e., 95.54 µm^3^ and 119 µm^2,^ respectively (see Table 1).

**Table 1:** Best shape parameters of a trophozoite-infected erythrocyte.

| Parameters | $D_0$ | $a_0$ | $a_1$ | $a_2$ |
| --- | --- | --- | --- | --- |
| Value | $7.0 \mu\text{m}$ | 0.132 | 2.026 | -4.491 |

### 2.2 Constitutive erythrocyte membrane model

The erythrocyte membrane constitutive model was developed based on the Yeoh strain energy density function [32–36] by incorporating the evolving membrane stiffness exponential term see Eqn. (2). The erythrocyte membrane is assumed to be homogeneous, isotropic, and nearly incompressible. The developed erythrocyte membrane model was calibrated by using the membrane stiffness parameter β_1_, ensuring that the predicted in-plane shear modulus, both at trophozoite and schizont stages, aligns with the previously determined values of 21.3 and 53.3 µN/m, for the trophozoite and schizont stages respectively [30]. The model was implemented in Abaqus utilizing a user material subroutine.

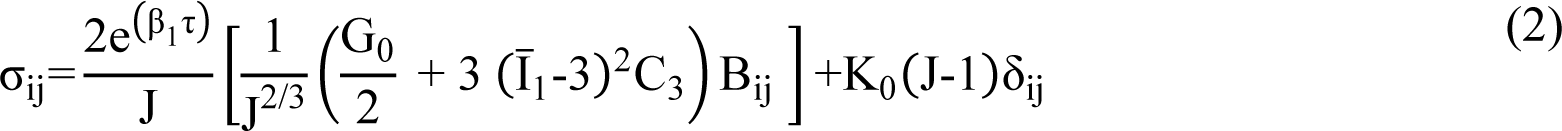

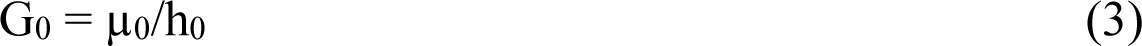

Where µ0 is the in-plane membrane shear modulus, h_0_ is the erythrocyte membrane thickness [37], C_3_ = G_0_/20 and K_0_ are the material parameters, J is the total volume ratio, τ is the simulation time, β_1_ is membrane stiffness parameter, ^I̅^_1_ is the first deviatoric strain invariant of the left Cauchy-Green deformation tensor B, see Eqn (4).

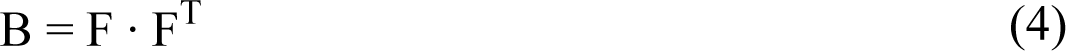

Where F is the deformation gradient tensor [38], and the invariant is defined as follows :

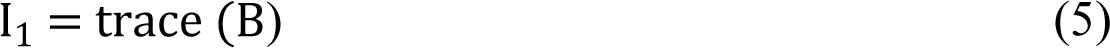

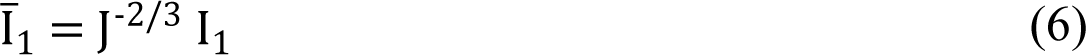

### 2.3 Modelling the erythrocyte cytoplasm

The erythrocyte cytoplasm is modelled as a homogeneous Newtonian fluid with a spherical void created for the spherical malaria parasite model. The smoothed particle hydrodynamics (SPH) method [39] was employed in Abaqus to discretize the cytoplasmic domain into SPH particles, facilitating its fluidization. This allows the cytoplasmic domain to undergo finite deformations induced by the expanding malaria parasite model.

### 2.4 The malaria parasite intra-erythrocyte growth model

The spherical malaria parasite FE model represents a single, nearly rigid merozoite that isotopically expands to mimic the volumetric growth of the malaria parasite due to schizogony. As such, the malaria parasite volumetric growth model was initialized when the volume of a sphere was 2.1 µm^3^, representing a 2.2% volume fraction (VF) of the malaria parasite in the infected erythrocyte. This volume is comparable to the previously determined value estimated from cryo x-ray images of a malaria merozoite where the maximum virtual volume, V_virtual,_ was estimated to be 2.06 µm^3^ [40]. Abaqus-defined isotropic thermal expansion model was employed and calibrated to emulate plasmodium volumetric growth due to schizogony. The thermal expansion model computes the strains by specifying the thermal load θ_i_ and the expansion coefficient α^i^, see Eqn (7).

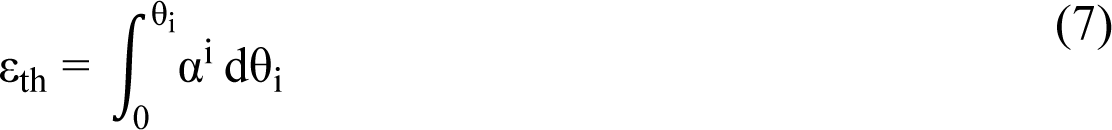

Where ε_th_ is the thermal strain α^i^ is the thermal coefficient of expansion, θ_i_ is the thermal load. The computed thermal strains are a driving force for volumetric growth, incrementally evolving the volume fraction (VF) of the malaria parasite model in the infected erythrocyte model from the point prior to the trophozoite stage to a point beyond the schizont stage. The malaria parasite growth model was iteratively calibrated by ensuring that the (VF) of malaria parasites in the infected erythrocyte evolves from 2.2 % (trophozoite VF) to 83.3 % (schizont VF prior to rupture). The metrics for calibration included: 1) material parameters for the spherical parasite model, i.e., linear elastic modulus (E), Poisson’s ratio (ν), and density. (ρ), 2) growth parameters for inducing volumetric expansion of the parasite domain, i.e., temperature load θ_i_ and the coefficient of expansion θ_i_. These calibration parameters (see Table 2) were exclusively utilized for volumetric prediction within the growth model, as the model interacts with other components, particularly the erythrocyte cytoplasm, volumetrically rather than thermally.

**Table 2:** Parameters for the Abaqus defined malaria parasite growth model.

| $\alpha^i$ | $\theta_i$ | N | E | P |
| --- | --- | --- | --- | --- |
| 0.8/°C | 20 °C | 0.3 | 1 kPa | 0.17 kg/m<sup>3</sup> |

### 2.5 Finite element modelling and analysis

The dynamic explicit analysis was utilized to compute the deformations in Abaqus. However, mass scaling was exclusively implemented on the erythrocyte membrane model, primarily aimed at obtaining a quasi-static response. The efficacy of the mass scaling algorithm in achieving a quasi-static solution was evaluated by ensuring that the total kinetic energy of the erythrocyte model remained significantly smaller than its internal or strain energy. In this context, kinetic energy accounts for inertia effects impacting the overall response of the erythrocyte model, while internal energy characterizes static effects. Time incrementation was automatically managed through built-in functionality. An adaptive, global estimation algorithm was employed to determine the maximum frequency of the entire model using the current dilatational wave speed.

The FE model comprises three components with various element lengths, i.e., the erythrocyte membrane, the spherical volume representing the parasites, and the cytoplasm, see Figure 1.

**Figure 1:**
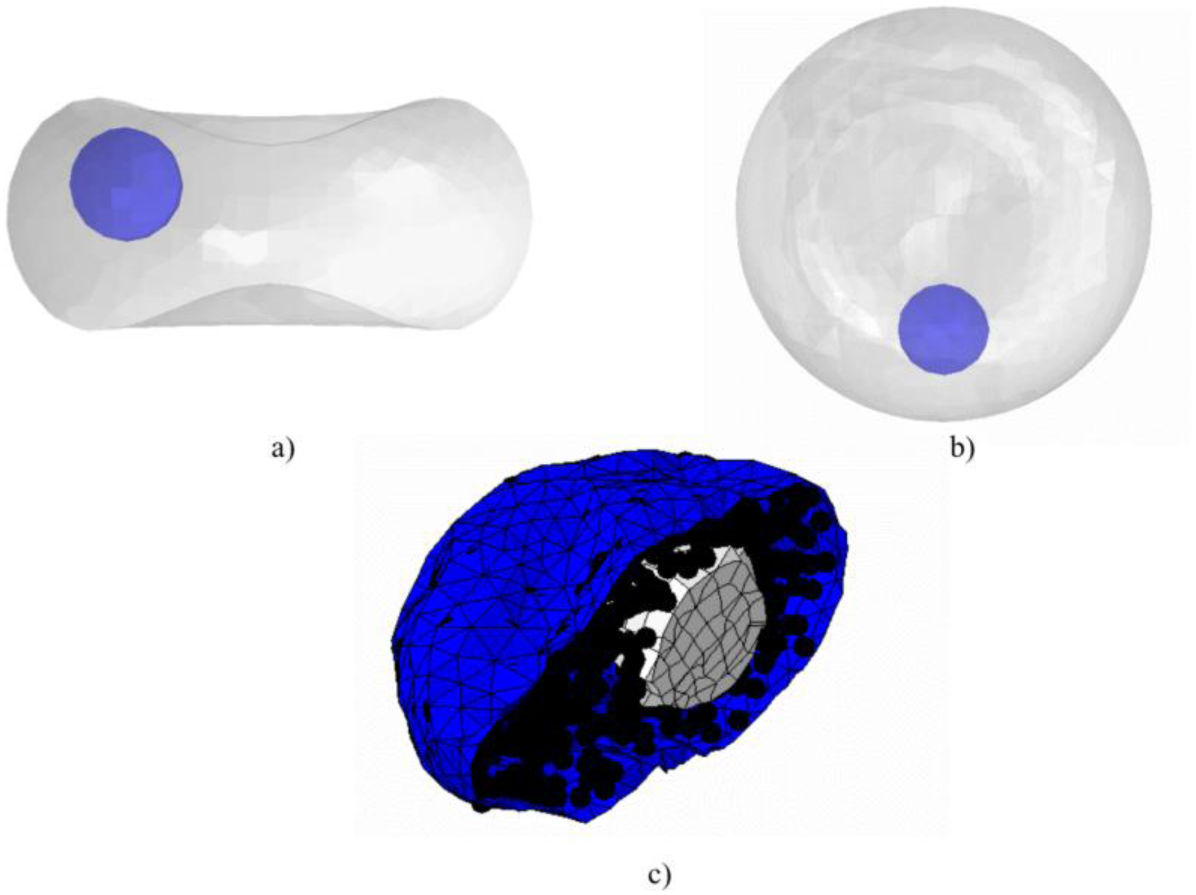
The malaria parasite spherical model (blue) of an infected erythrocyte (a) and (b), erythrocyte membrane (blue) meshed with triangular shell elements, erythrocyte cytoplasm SPH particles (black) generated from the 8-node linear brick elements with reduced integration and hourglass control (C3D8R) (b), the expanded malaria parasite spherical model (white) meshed using a ten-node modified quadratic tetrahedron elements (C3D10M) (c).

**Figure 2:**
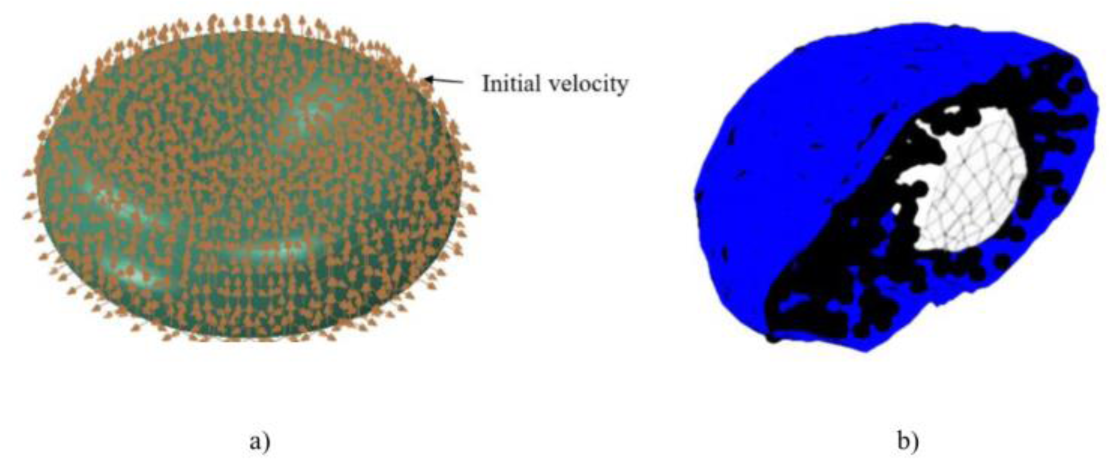
The initial velocity of 2×10^-30^ mm/s applied at each node of the erythrocyte membrane (a), the malaria parasite model with growth (b).

**Figure 3:**
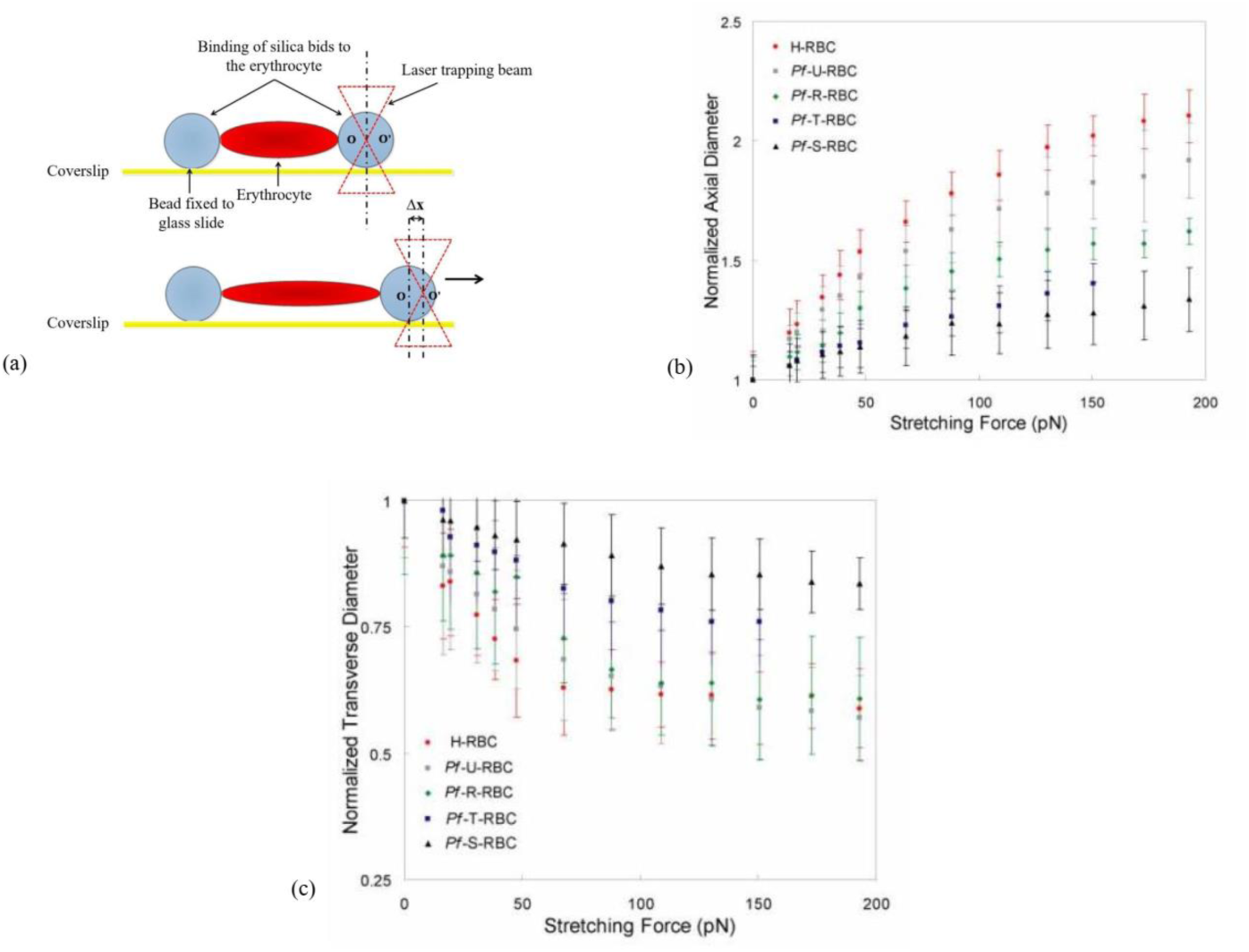
Optical tweezer experiment set up (a), normalized erythrocyte axial diameter (b), normalized erythrocyte transverse diameter (c), sited with permission from [30, 42].

In Abaqus, the erythrocyte membrane was modelled using shell elements with a thickness of 0.01 µm [41]. SPH algorithm was applied only to the erythrocyte cytoplasmic domain. Three-node triangular shell elements with reduced time integration (S3R) were used to mesh the erythrocyte membrane. The reduced time integration algorithm provided more accurate results and reduced the running time. The erythrocyte cytoplasm was meshed using the 8-node linear brick elements with reduced integration and hourglass control (C3D8R). The mesh provides the initial spatial particle discretization required for the SPH algorithm, while the ten-node modified quadratic tetrahedron elements (C3D10M) mesh the malaria parasite model. The details of the mesh sizes are provided in Table 3 below.

**Table 3:** Element type and mesh sizes for various components of the infected erythrocyte model.

| Structure | Element type | Elements |
| --- | --- | --- |
| Membrane | S3R | 19942 |
| Cytoplasm | C3D8R | 841 |
| Parasite model | C3D10M | 700 |

**Table 4:**
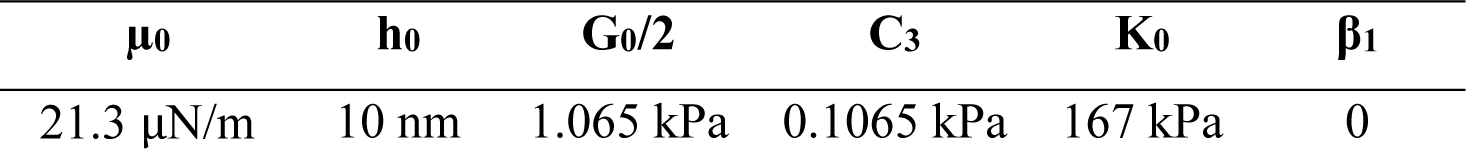
Best fit material parameters for the erythrocyte membrane

| $\mu_0$ | $h_0$ | $G_0/2$ | $C_3$ | $K_0$ | $\beta_1$ |
| --- | --- | --- | --- | --- | --- |
| 21.3 $\mu\text{N/m}$ | 10 nm | 1.065 kPa | 0.1065 kPa | 167 kPa | 0 |

**Table 5:** Best fit shape parameters for the erythrocyte, the surface area to volume ratio (S/V) of a healthy and trophozoite infected erythrocyte

| Erythrocyte shape | Surface area (S) | Volume (V) | S/V |
| --- | --- | --- | --- |
| Healthy | 135 $\mu\text{m}^2$ | 94 $\mu\text{m}^3$ | 1.44/$\mu\text{m}$ |
| T-Infected | 119 $\mu\text{m}^2$ | 95.54 $\mu\text{m}^3$ | 1.05/$\mu\text{m}$ |

The interaction module in Abaqus Explicit was employed to define the mechanical interaction of the interfaces of structures involved during the malaria parasite intra-erythrocyte development stages. The general contact algorithm allowed elementary contact definitions with few interface restrictions.

The deformation of the developed infected erythrocyte model at the trophozoite stage, predicted computationally from the optical tweezer simulation experiment implemented in Abaqus, was compared against previously determined data from an optical tweezer experiment [30], where a 200 pN force was applied to stretch the trophozoite infected erythrocyte.

## 3 Results

### 3.1 Validation of the malaria parasite-infected erythrocyte finite element model

The developed trophozoite-infected erythrocyte model was validated by comparing optical tweezer experimental data [30] with the numerical data obtained by simulating an optical tweezer experiment. When a force of 0.21 nN (see Figure 4a) is applied diametrically, the diameter of the infected erythrocyte FE model increases axially in the direction of the applied load (see Figure 4b) and decreases transversely in the direction normal to the applied force (see Figure 4c). The diameter of the infected erythrocyte FE model that increases in the applied load direction is referred to as axial diameter, whereas the diameter of the infected erythrocyte FE model that decreases in the direction normal to the applied force is referred to as the transverse diameter.

**Figure 4:**
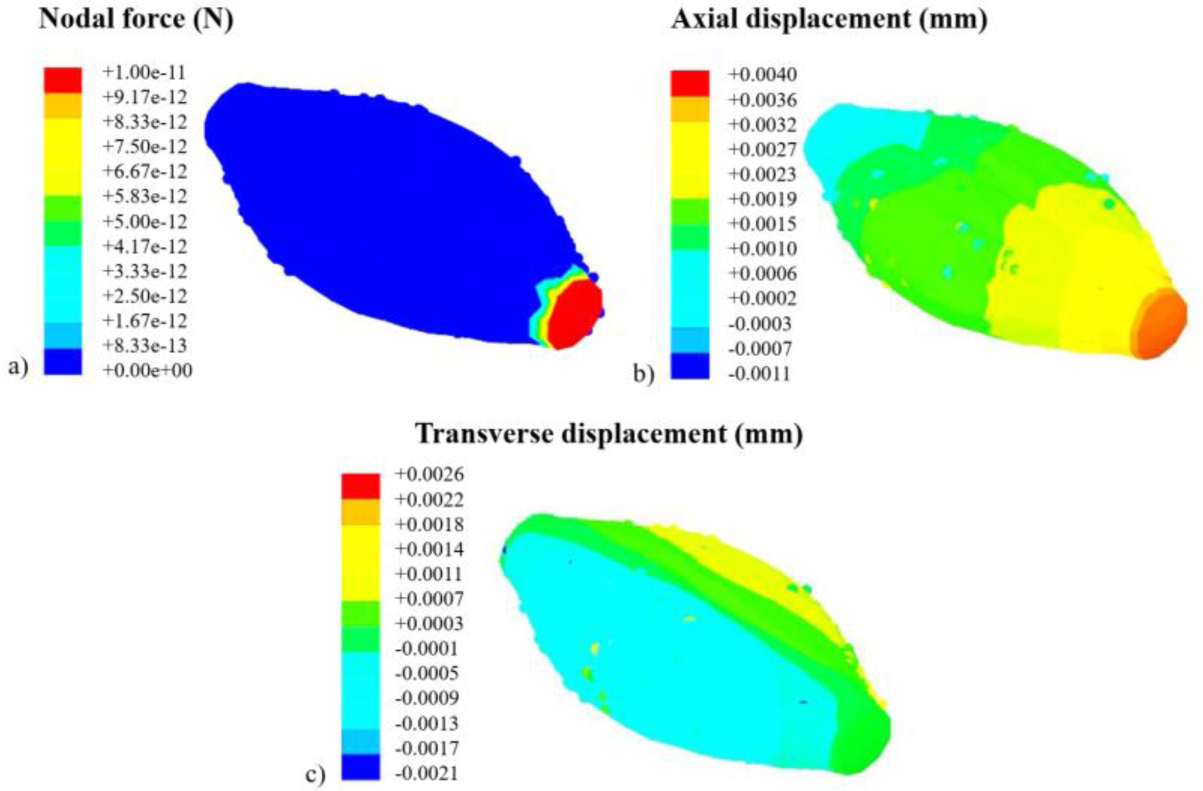
The contour plot of the erythrocyte (a) nodal force, where the total force for 20 nodes represented by the red contours is equal to 0.21 nN, (b) axial displacements, and (c) transverse displacement. [30].

The normalized axial and transverse diameters of the infected erythrocyte model determined from the optical tweezer simulation align with experimental data (see Figure 5). The predicted normalized axial and transverse diameters of 1.47 and 0.85 correspond to the maximum applied stretching force of 0.21 nN.

**Figure 5:**
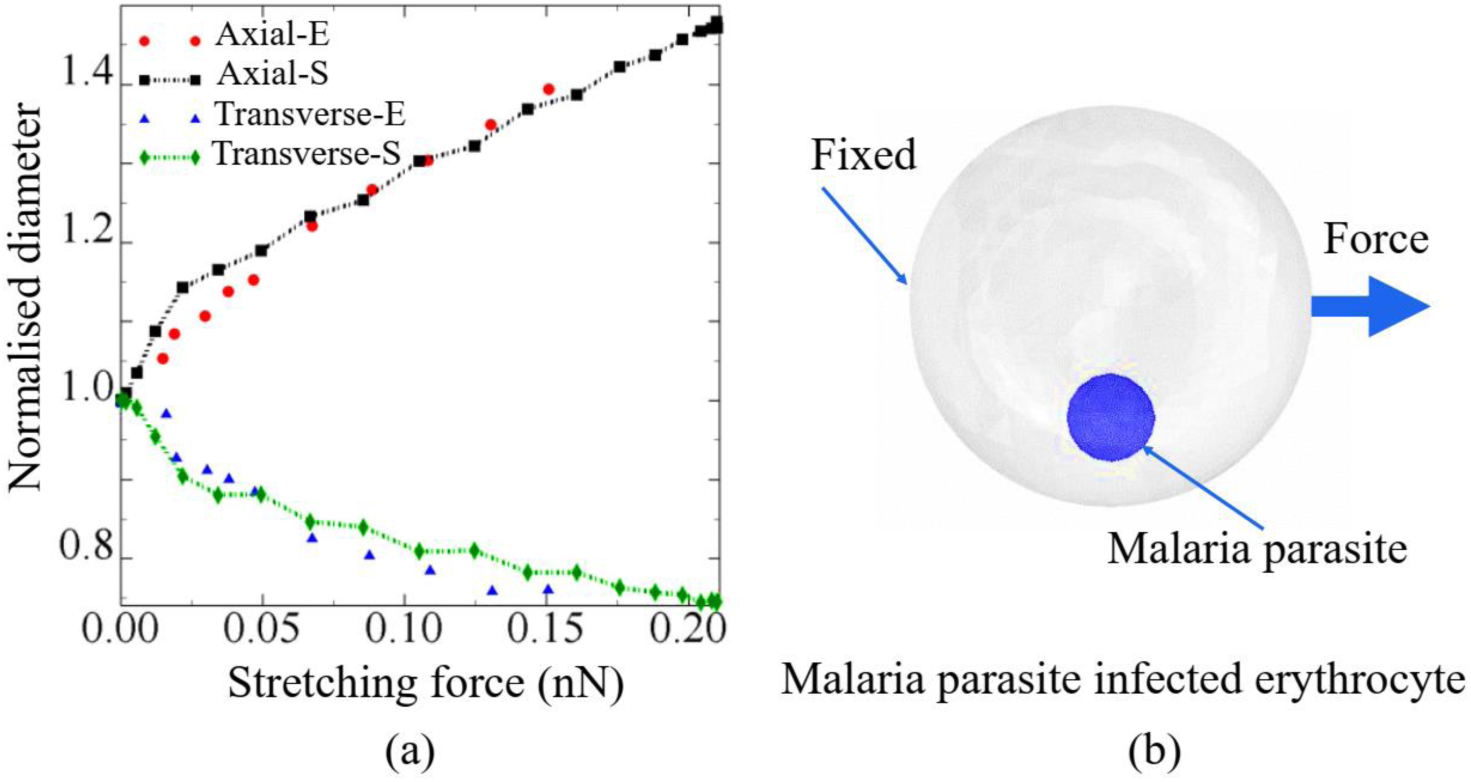
A comparison between optical tweezer simulation data (axial and transverse normalized erythrocyte diameters) and previous optical tweezer data (axial and transverse diameters of the trophozoite-infected erythrocyte) [30].

### 3.2 Model calibration

The membrane stiffness parameter was employed to iteratively calibrate the developed membrane model by altering β_1_ between 0.49 and 2 and comparing the minimum and maximum predicted in-plane shear modulus values with the previously determined values of the erythrocyte membrane at trophozoite and schizont stages, respectively. When β_1_ was equal to 0.49, the in-plane erythrocyte membrane shear modulus evolved from 21.8 µN/m to 35.69 µN/m whereas the parasite volume fraction in the infected erythrocyte evolved from 2.2 % to 107%, (see Figure 6a). Similarly, for β_1_ = 1, the in-plane erythrocyte membrane shear modulus evolved from 22.5 µN/m to 61.8 µN/m while the malaria parasite volume fraction evolved from 2.2% to 104% (see Figure 6). The malaria parasite growth model was calibrated utilizing the thermal expansion coefficient α and thermal load. When α = 0.8/℃ and θ = 20 ℃, the malaria parasite volume fraction in the infected erythrocyte evolves from 2.2 % to 104% and107%, depending on the value of β_1_ (see Figure 6).

**Figure 6:**
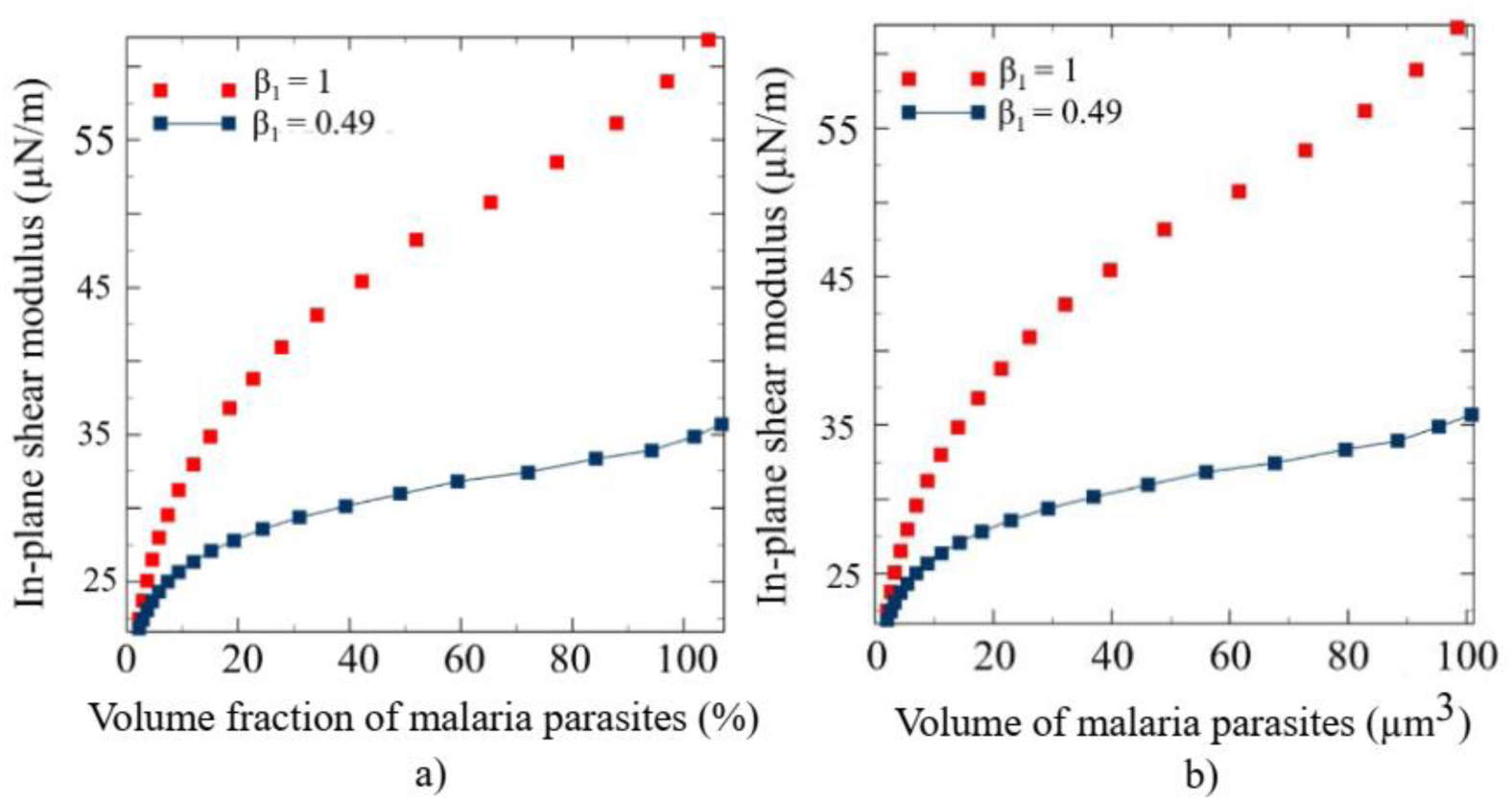
In-plane shear modulus of the erythrocyte membrane vs volume fraction of malaria parasites (a), and the in-plane shear modulus of the erythrocyte membrane vs volume of malaria parasites (b).

### 3.3 A mechanistic link between the parasite growth, malaria-induced erythrocyte membrane remodelling, and erythrocyte membrane areal strain

When the erythrocyte membrane stiffness parameter β_1_ is equal to 1, the developed model predicts the erythrocyte in-plane shear modulus of 51.8 µN/m and 57.5 µN/m corresponding to established the lower and upper bounds of erythrocyte lysis areal strain of 2 % and 4 % respectively and while the corresponding predicted volume fractions of malaria parasites in infected erythrocyte are 68.7% and 90 % respectively (see Figure 7a). The infected erythrocyte ruptures when the volume fraction of malaria parasites in the infected erythrocyte is more than 80% during the late schizont stage [11, 14, 30, 31]. At the known rupture volume fraction of the malaria parasite of 83 % (see Figure 8c), the predicted membrane areal strain and in-plane shear modulus are 3.1 % and 55.8 µN/m, respectively (see Figure 7a). Similarly, for β_1_ = 2, the predicted in-plane shear modulus are 107 µN/m and 147 µN/m at the lower and upper bound of erythrocyte membrane lysis erythrocyte areal strain, respectively, while the corresponding volumetric fractions of malaria parasites in infected erythrocyte are at 55.8 % and 92.4 % respectively (see Figure 7a). Additionally, at 83 % volume fraction, the erythrocyte membrane areal strain and the in-plane shear modulus are 3.2% and 131 µN/m, respectively.

**Figure 7:**
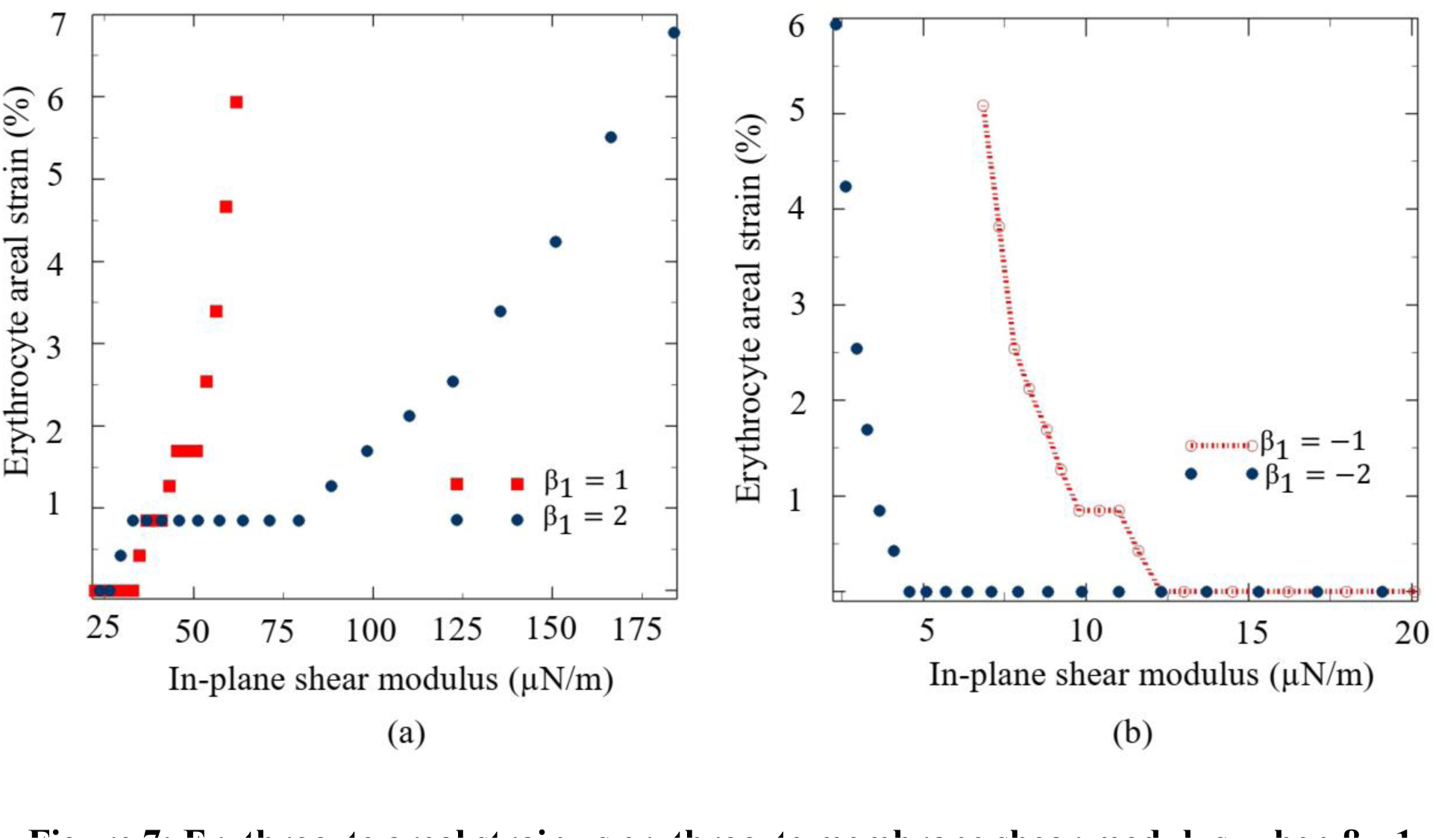
Erythrocyte areal strain vs erythrocyte membrane shear modulus, when β_1_=1 and 2 (a), when β_1_= -1 and -2 (b).

**Figure 8:**
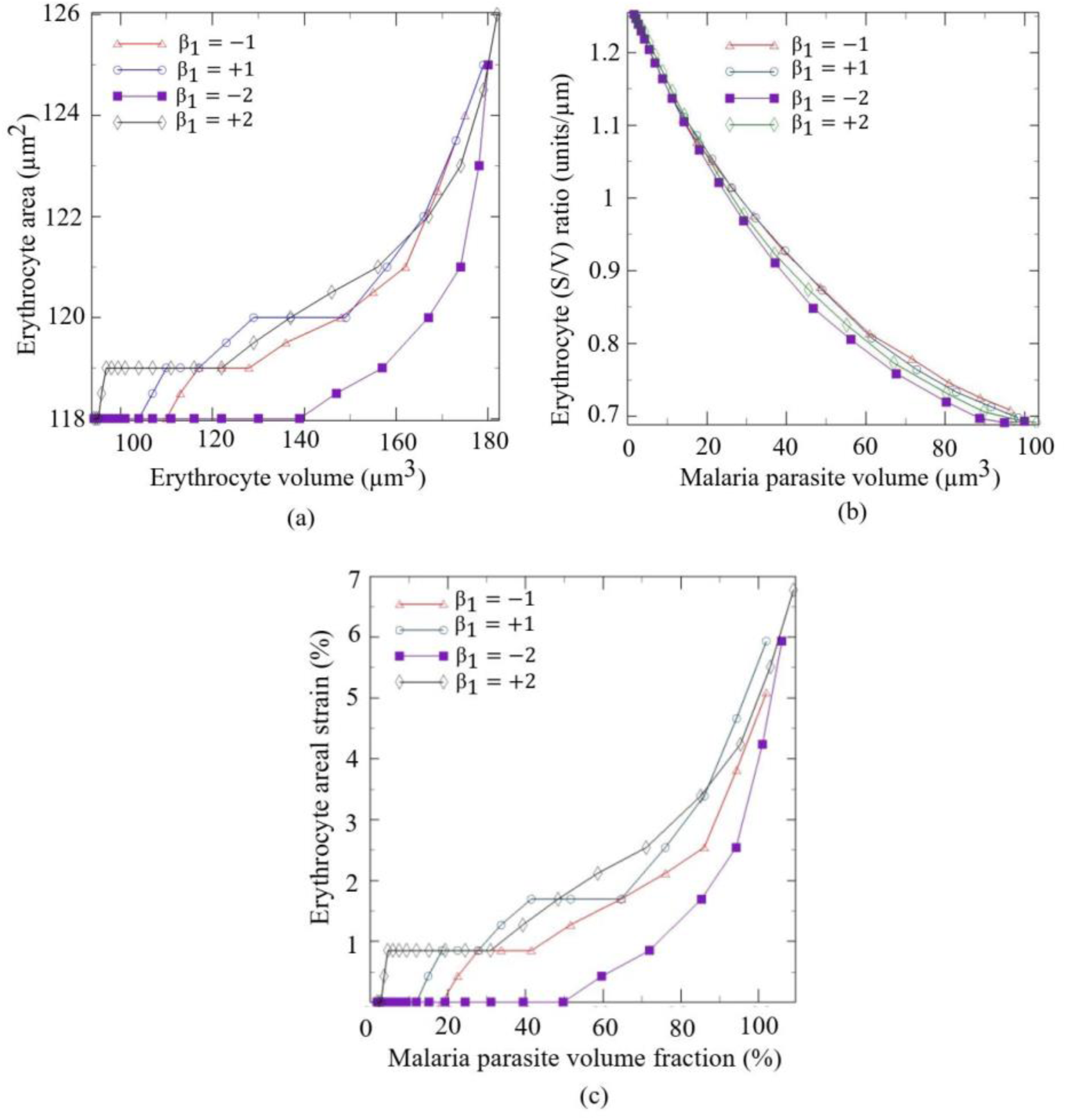
Erythrocyte areal strain vs erythrocyte volume (a), erythrocyte S/V vs malaria parasites volume (b), erythrocyte areal strain vs malaria parasites volume fraction in the erythrocyte.

In contrast to evaluating the impact of increasing the erythrocyte membrane in-plane shear modulus, the developed model was also utilized to explore the impact of reducing the erythrocyte membrane in-plane shear modulus on the infected erythrocyte rupture threshold using erythrocyte lysis areal strain as a rupture metric. Similarly, for β_1_ = -1, the predicted erythrocyte volume fraction of malaria parasites in infected erythrocytes and the in-plane shear modulus at the lower bound of erythrocyte membrane lysis are 72.8 %, and 8.4 µN/m respectively. Whereas at the upper bound of erythrocyte membrane lysis, when the erythrocyte areal strain is 4%, the volume fraction of malaria parasites in infected erythrocyte, and the in-plane shear modulus are 95.5 %, and 7.26 µN/m respectively (see Figure 7b and Figure 8c). The in-plane erythrocyte membrane shear modulus was further reduced by setting β_1_ to -2 to evaluate the implications of severely reducing the erythrocyte membrane shear modulus on the erythrocyte areal strain. The aim was to establish the correlation between reduced erythrocyte membrane stiffness and the timing of erythrocyte rupture, whether occurring early or late in the rupture process. At the lower bound of erythrocyte membrane lysis, when the erythrocyte areal strain is 2 %, the corresponding volume fraction of malaria parasites in infected erythrocyte, and the in-plane shear modulus are 88.5 %, for β_1_ = -2 (see Figure 8c) and 3.1 µN/m respectively (see Figure 7b) whereas at the upper bound of erythrocyte membrane lysis when the erythrocyte areal strain is 4 %, the predicted in-plane shear modulus is 2.66 µN/m (see Figure 7b), while the volume fraction of malaria parasites in infected erythrocyte is 99.8 % (see Figure 8c).

For the selected values of membrane stiffness parameter (β_1_=1,-1,2,-2), the infected erythrocyte surface area increases with respect to its volume; however, for severely softened erythrocyte membrane, i.e., when β_1_=-2, the surface area only rapidly changes when the erythrocyte volume is above 139 µm^3^ (see Figure 8a). The erythrocyte surface area to volume ratio depends on the relative increase of the erythrocyte surface area with respect to the erythrocyte volume. The erythrocyte surface area to volume ratio (S/V) decreases from 1.252 µm^-1^ to 0.693 µm^-1^, 1.252 µm^-1^ to 0.691 µm^-1^, for (β_1_=-2) and (β_1_=+2) respectively, whereas for (β_1_=-1) and (β_1_=+1), the erythrocyte surface area to volume ratio (S/V) decreases from (1.252 µm^-1^ to 0.708 µm^-1^) and (from1.252 µm^-1^ to 0.698 µm^-1^) respectively (see Figure 8b).

## 4 Discussion

Previous studies have achieved tangible progress in characterising biological factors that determine the exit of the malaria parasites from infected erythrocytes during the late schizont stage. However, many of these studies do not address the role played by malaria parasite-mediated mechanical deformations on eryptosis [43, 44]. To date, mechanistic insights describing the connection between eryptosis and the mechanics of the infected erythrocyte during the growth and exit of malaria parasites remain limited. This is due to the difficulty of conducting experimental studies on eryptosis involving malaria infection. This is mainly because eryptosis read-outs in vitro studies primarily depend on culture conditions. In addition, the conditions often used to study eryptosis do not support plasmodium falciparum growth and are unable to prime erythrocytes for eryptosis [6]. Computational studies, on the contrary, provide an easier alternative approach for investigating the exit of malaria parasites from the infected erythrocyte. The current research aimed to develop an in-silico model for investigating the impact of malaria parasite-induced erythrocyte morphology changes and malaria parasite-induced remodelling of erythrocyte membrane on the eryptosis.

### 4.1 Validation of the trophozoite-infected erythrocyte model

The developed malaria parasite-infected erythrocyte model was validated at 2.2 % malaria parasite volume fraction, representing an early stage of malaria parasite infection by comparing the deformations of the erythrocyte model computed from the optical tweezer simulation with the previously determined optical tweezer experimental data (see Figure 5) [30, 36].

Interestingly, the diametric numerical data computed from the developed malaria parasite erythrocyte model aligns with diametric data from the optical tweezer experiment [30, 36] (see Figure 5).

### 4.2 Validation of the malaria parasite growth and erythrocyte membrane model at the trophozoite and schizont stage

The malaria parasite volume fraction and the erythrocyte membrane in-plane shear modulus were used as metrics for calibrating both the parasite growth and the developed erythrocyte membrane model. Parasite growth and erythrocyte membrane remodelling were concurrently induced from the trophozoite stage (VF = 2.2 %) to a point slightly beyond the schizont stage (VF = 102 %).

The models were calibrated at both the trophozoite and schizont stages by comparing both the volume fractions of the malaria parasite in the infected erythrocyte and the in-plane shear modulus of the erythrocyte membrane with previously determined data. The rationale for this comparison was to validate the accuracy of the predictions for both the parasite growth and erythrocyte membrane model since, from the trophozoite to the schizont stage, the erythrocyte membrane properties evolve due to remodelling, while the malaria parasites undergo collective volumetric expansion due to schizogony. The minimum volume of the malaria parasites at the trophozoite stage was assumed to be 2.1 µm^3^ (VF = 2.2 %), representing an idealized virtual volume of a single merozoite of 2.06 µm^3^ [40]. Whereas the maximum total volume of the malaria parasites at the end of the schizont stage, prior to rupture, is estimated as 63 µm^3^, representing an 83% volume fraction of the malaria parasites in the infected erythrocyte [11]. These values along with the average previously determined shear modulus values of the erythrocyte membrane at the trophozoite and schizont stages of 21.3 and 53.3 µN/m, respectively, closely align with the predictions of the developed model when β_1_=1. The predicted in-plane shear modulus of the erythrocyte membrane is 55.3 µN/m approaching the previously predicted average value of 53.3 µN/m [30], with an error of 3.8 % at the volume fraction of 83% prior to rupture (see Figure 6), and when the areal strain of the erythrocyte membrane is 3.1%.

Similarly, the predicted in-plane shear modulus of the erythrocyte membrane at the trophozoite stage is 22.5 µN/m and also nearing the previously predicted average value of 21.3 µN/m [30] with an error of 5 %. These results demonstrate the accuracy of the malaria parasite growth model as well as the accuracy of the erythrocyte membrane model. In addition, the developed model reveals that the rupture mechanism is associated with erythrocyte membrane areal strain-induced lysis. i.e., when exceeded cell lysis occurs.

### 4.3 Mechanistic impact of the malaria parasite volume fraction and erythrocyte membrane in-plane shear modulus on the erythrocyte membrane areal strain

The developed and validated model was utilized to establish correlations between the volume fraction of malaria parasites in the infected erythrocyte, the in-plane shear modulus of the erythrocyte membrane, and the lysis areal strains. The infected erythrocyte ruptures when the volume fraction of malaria parasites in the infected erythrocyte is more than 80% during the late schizont stage [31]. Additionally, it is well known that the erythrocyte membrane undergoes lysis, referred to as eryptosis when the erythrocyte membrane areal strain of 2-4% is exceeded [21], it is therefore apparent that the increase in parasite volume in the infected erythrocyte leads to the increase in the erythrocyte membrane areal strain, a metric for cell lysis. It is well known that the malaria parasite induces remodelling of the erythrocyte membrane to stiffen the erythrocyte membrane. However, to date, how the remodelling of the erythrocyte membrane correlates to the areal strain induced through malaria parasite volumetric growth remains unknown. Furthermore, whether the rupture process of the erythrocyte membrane is linked to lysis areal strains remains a mystery. The result from the developed model sheds some light on the effect of remodelling on the areal strain. Our results demonstrate that severely altering the mechanical properties of the erythrocyte membrane can affect the timing of rupture, i.e., for β_1_ = 2, the erythrocyte membrane areal strain first changes from zero to 0.424 % much earlier when the malaria parasite volume fraction (VF) is 0.358 %, followed by similar instances when β_1_=1 occurring at VF = 14.9 %, when β_1_=-1 occurring at VF= 22.5 %, and when β_1_ =-2 occurring at VF = 59.6%, (Figure 8c). Additionally, at the established rupture volume fraction of 83% when β_1_=1, the corresponding erythrocyte membrane areal strain of 3.1% is lower than the areal strain of 2.4 %., when β_1_=-1. Similarly, as the stiffness parameter β_1_ decreases with a unit decrement from 2 to -2, the areal strain decreases from 3.2 to 1.55, at the established rupture volume fraction of 83%, implying that decreasing the stiffness of erythrocyte membrane during the malaria intra-erythrocytic development stage can potentially reduce the erythrocyte membrane areal strain preventing the lysis areal strain from being attained. In contrast, increasing the erythrocyte membrane stiffness increases the areal strain, allowing lysis-areal strain to be attained. Inducing early erythrocyte rupture can be useful in minimizing the virulence of malaria infection by inducing rupture to expose the malaria parasite before they have excessively replicated to overwhelm the human host’s immune system. However, further experiment studies need to be conducted to validate this finding of whether strain hardening of the erythrocyte membrane induces early rupture, and whether can this be a potential therapeutic intervention to minimize the virulence of the malaria parasites.

### 4.4 Rupture in-plane shear modulus of the erythrocyte membrane falls within the predicted lysis shear modulus range when β1 = 1

When the erythrocyte membrane stiffness parameter β_1_ is equal to 1, the developed model predicts the erythrocyte in-plane shear modulus of 51.8 µN/m and 57.5 µN/m, corresponding to the lower and upper bounds of erythrocyte lysis areal strain of 2 % and 4 % respectively, corresponding to predicted volume fractions of malaria parasites in infected erythrocyte are of 68.7% and 90 % respectively (see Figure 7a). Interestingly, the previously predicted rupture shear modulus of 53.3 µN/m [30] and the previously determined malaria parasite volumetric threshold for erythrocyte rupture above 80 % aligns with the previously predicted range. The result from the developed model aligns with the previously determined thresholds for rupture of the infected erythrocyte and demonstrates the accuracy of the predictions of the developed model.

### 4.5 Impact membrane shear modulus on rupture thresholds

In contrast, we explore the consequences of reducing the erythrocyte membrane in-plane shear modulus (β_1_ = -1 and -2) on the infected erythrocyte rupture threshold and lowering the shear modulus to 8.4 µN/m and 3.1 µN/m at the lower bound of lysis results in predicted volume fractions of 72.8% and 88.5%, respectively. At the upper bound (4% erythrocyte areal strain), the volume fractions increase to 95.5% and 99.8%, illustrating the sensitivity of rupture thresholds to membrane softening (see Figure 7b and Figure 8c). At the lower bound of the erythrocyte membrane lysis, for β_1_=-2, the volume fraction of the malaria parasites in the infected erythrocyte is 88.5% higher than the rapture erythrocyte membrane threshold. Whereas at the upper bound of erythrocyte membrane lysis, the volume fraction of the malaria parasite in the infected erythrocyte is 99.8% greater than the rupture erythrocyte membrane threshold.

### 4.6 Surface area to volume ratio dynamics

The erythrocyte surface area to volume ratio (S/V) [28] exhibits noteworthy variations under different membrane stiffness conditions. For severely softened membranes (β1=-2), the S/V ratio experiences a rapid change only when the erythrocyte volume exceeds 140 µm³. This indicates the critical influence of membrane stiffness on the deformability of infected erythrocytes. Hence influencing the timing of erythrocyte rupture.

## 5 Conclusion

In summary, our investigation underscores the intricate interplay between membrane properties, parasite volume fractions, and erythrocyte rupture thresholds, providing valuable insights into the mechanics of malaria-induced erythrocyte remodelling. This understanding holds significant implications for therapeutic interventions and is a foundation for further investigations into erytosis.

The developed model has played a crucial role in analyzing the mechanics involving the exit of malaria parasites from infected erythrocytes, with a specific focus on factors such as erythrocyte membrane stiffness and intra-erythrocyte replication of malaria parasites. The developed finite element analysis model has elucidated the relationship between erythrocyte membrane stiffness, parasite volume fractions, and erythrocyte rupture thresholds, offering valuable insights into the mechanics of malaria-induced erythrocyte remodelling and its potential application in predicting the timing of parasite egress. Notably, the developed model reveals that the rupture of the erythrocyte membrane occurs within the range of 2-4% areal strain, aligning with the range associated with erythrocyte lysis. This emphasizes the potential role of erythrocyte membrane areal strain in driving eryptosis during malaria infection. Understanding the growth and exit mechanics of malaria parasites provides insights into their survival mechanism within the human host offering potential antimalarial targets.

## Funding

This research was supported financially by the Malawi University of Business and Applied Sciences. The funder had no role in study design, data collection and analysis, decision to publish, or manuscript preparation. Any opinions, findings, conclusions, or recommendations expressed in this publication are those of the authors and do not represent the official views of the funding agency.

### Authorship contribution statement

Chimwemwe Msosa: Writing - review & editing, Writing – original draft, Visualization, Validation, Software, Project administration, Methodology, Investigation, Funding acquisition, Formal analysis, Data curation, Conceptualization.

YD Motchon: Writing - review & editing, Visualization, Validation, Software, Methodology, Investigation.

### Data availability

Data supporting the results of this article are available on: https://doi.org/10.6084/m9.figshare.25833301.v2

## Nomenclature

a_0_: Erythrocyte shape parameter
a_1_: Erythrocyte shape parameter
a_2_: Erythrocyte shape parameter
D_0_: Erythrocyte diameter
ε_th_: Thermal strain
θ: Thermal load
α: Coefficient of expansion
J: Total volume ratio
τ: Simulation time
B: Left Cauchy deformation tensor
µ_0_: In-plane shear modulus
C_3_: Material parameter
C3D10M: Ten-node modified quadratic tetrahedron elements
C3D8R: 8-node linear brick elements with reduced integration and hourglass control
E: Elastic modulus
F: Deformation gradient tensor
G_0_: Shear modulus
h_0_: The thickness of the erythrocyte membrane
I_1_: First strain invariant
K_0_: Initial bulk modulus
S3R: Triangular shell elements with reduced time integration
X: x coordinate point on the surface of the erythrocyte
Y: y coordinate point on the surface of the erythrocyte
Z: z coordinate point on the surface of the erythrocyte
β_1_: Membrane stiffness parameter
δ: Kronecker delta function

